# scJET: Full-gene Space Single-cell Expression Generation with Patch-based Transformer Modeling

**DOI:** 10.64898/2026.07.06.736701

**Authors:** Qiantong Liang, Qing Rex Lyu

## Abstract

Most single-cell generative models rely on highly variable genes (HVGs) or low-dimensional latent representations, limiting their capacity to capture the complexity of full-gene features. We present scJET, a patch-based Transformer denoising framework that operates in full-gene space. scJET preserves global manifold structure, local neighborhood statistics, and gene-level expression programs. By combining scalable patch tokenization with full-gene denoising, scJET provides an efficient framework for transcriptome-wide single-cell matrix generation.

---

Artificial-intelligence virtual cells (AIVCs) require models that extend experimental single-cell measurements beyond the finite conditions sampled in the laboratory [1–3]. Such models must represent unobserved or sparsely sampled cell states, continuous transitions, missing expression patterns, and transcriptome-wide gene programs. Single-cell generative models address this need by learning expression distributions and generating or reconstructing plausible single-cell expression matrices, providing a computational foundation for transcriptome-wide modeling [4–8].

Existing single-cell generative models often improve scalability by selecting highly variable genes (HVGs) or learning low-dimensional latent representations [9–12]. Although efficient, these strategies can restrict the output space and reduce direct access to genes that are lowly variable, context-specific, or functionally relevant. HVG-based feature selection can omit low-variance functional genes, whereas latent representations can compress the gene expression structure and reduce interpretability. These limitations are particularly relevant for AIVCs, which require comprehensive transcriptome-wide representations of cellular states rather than profiles restricted to selected features or compressed embeddings. Thus, generation of single-cell matrices for full-gene space offers a way to preserve transcriptome-wide information and may help model gene-level expression relationships that are lost under HVG-based feature selection or latent compression. This motivates generation directly in full-gene space to preserve transcriptome structure.

Current diffusion models typically predict noise or noisy quantities rather than clean data directly [13–15]. Recently, Li and He argue that these two objectives are fundamentally different because natural data lie on a low-dimensional manifold, whereas noisy quantities lack this structure [16]. Based on this idea, JiT (Just Image Transformer) uses large-patch Transformers directly on raw pixels to predict clean data, requiring no tokenizer, pretraining, or extra loss, and achieves competitive generative performance [16]. We used this clean-data prediction principle as an algorithmic motivation and adapted it to single-cell expression matrices.

Here, we present scJET (single-cell Just Expression Transformer), a patched Transformer-based denoising framework that operates directly in full-gene expression space. scJET divides each expression profile into expression patches and uses them as computational tokens (Fig. 1a). scJET represents expression profiles as vector patches for efficient Transformer modeling and learns to recover clean expression states from corrupted inputs, enabling unified unconditional, conditional, and trajectory-based generation without latent representation or feature selection (Fig. 1c).

**Fig. 1.**
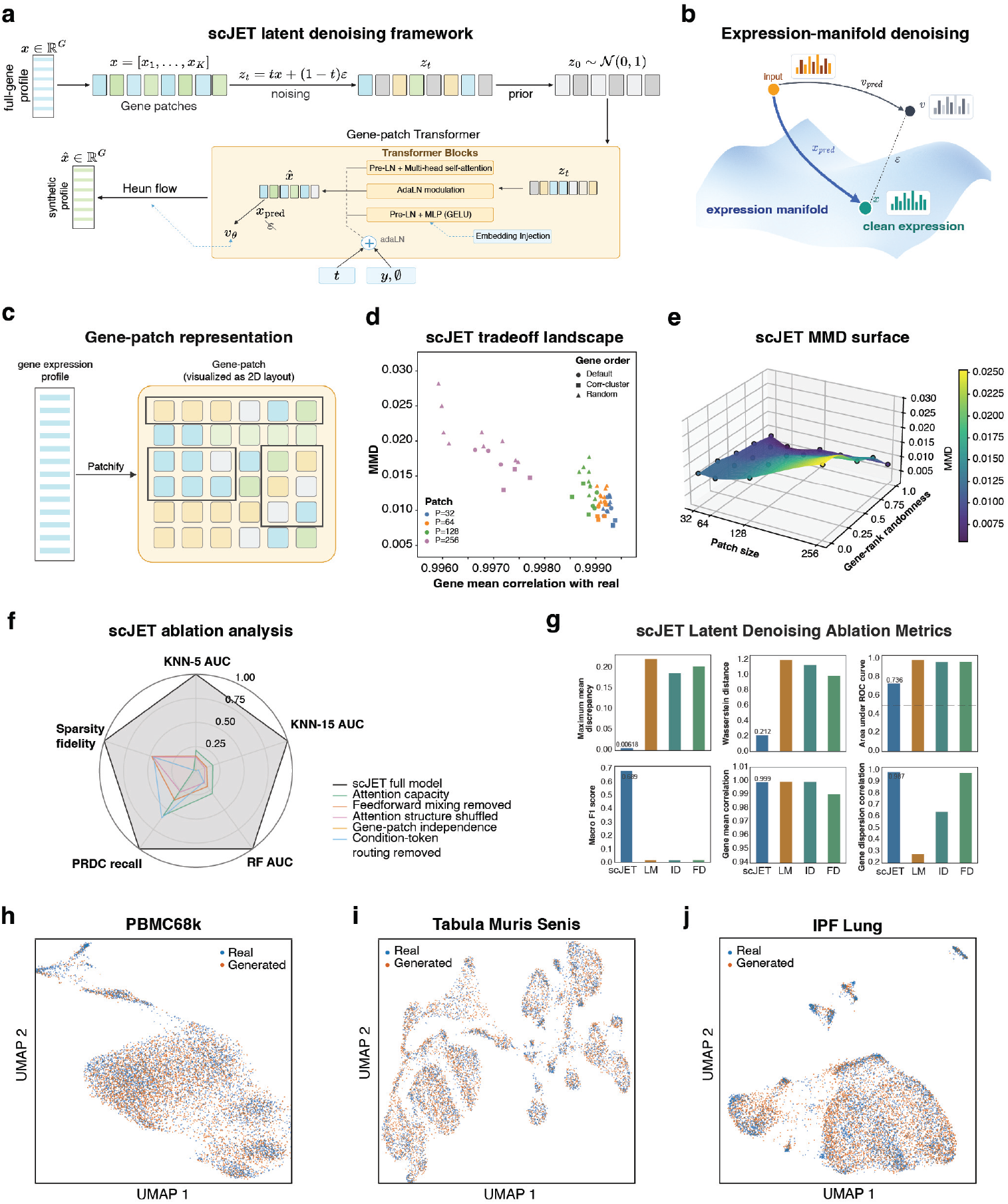
scJET framework for full-gene single-cell expression modeling. (a) Overview of scJET framework. (b) Expression-manifold denoising interpretation. (c) Gene-patch representation of full-gene expression profiles. (d) Tradeoff landscape between gene-level correlation and distributional discrepancy (MMD) under different patch sizes and gene ordering strategies. (e) MMD surface over patch size and gene-order perturbations. (f) Ablation analysis of scJET. (g) Comparison between scJET and latent/decoder-based variants. (h–j) Unconditional generation on PBMC68k, Tabula Muris Senis and IPF Lung datasets.

We formulate single-cell generation as denoising on a structured high-dimensional expression manifold (Fig. 1b). We assume that cellular transcriptomes reside on a biological manifold shaped by cell identity, lineage, and functional state, while corrupted samples deviate from it [17].

Let *x* ∈ ℝ^*G*^ denote a standardized full-gene expression profile. We construct a corrupted state *z*_*t*_ by mixing *x* with Gaussian noise over continuous time *t* ∈ (0, 1). scJET learns a denoising function

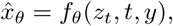

where *y* represents optional biological conditions, such as cell type, tissue, or experimental state. The input *z*_*t*_ is partitioned into expression patches, embedded as tokens, and combined with denoising-time and condition embeddings. The Transformer models dependencies across patches and directly predicts the clean expression target 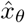 in the original *G*-dimensional gene space. Thus, scJET differs from latent or noise-prediction models and maintains gene-level coordinates throughout generation. During sampling, noise is iteratively removed to recover the learned manifold, producing full-gene profiles after inverse normalization.

Because genes do not have a natural sequential ordering, we tested sensitivity to gene patching and ordering. We varied patch size and reordered genes using default, randomized, and correlation-clustered arrangements (Fig. 1d, Extended Data Fig. 1a,b). scJET remained stable across perturbations, maintaining strong gene-level correlation and low MMD. Gene ordering caused only minor variation within patch regimes, indicating limited dependence on local adjacency. Consistently, continuous gene-rank perturbation produced smooth MMD changes (Fig. 1e). Thus, patch size affects resolution, but local gene grouping is not critical. These results indicate that scJET relies primarily on global expression structure.

We next evaluated architectural components. Full scJET consistently outperformed ablated variants across kNN distinguishability, RF classification, PRDC recall, and sparsity fidelity (Fig. 1f). All architectural modifications—including reducing attention capacity, removing feed-forward mixing, shuffling attention structure, enforcing gene-patch independence, or removing condition-token routing—consistently degraded performance. We also compared full-gene scJET with latent or decoder-based denoising variants. Although some latent variants retained high gene-mean agreement, they exhibited substantially larger MMD and Wasserstein distance, higher real–generated discrimination AUC, weaker condition fidelity, and reduced gene-dispersion agreement (Fig. 1g). Overall, these ablations support the importance of scJET’s decoder-free, full-gene, clean-target denoising design.

To evaluate scJET across complementary biological and statistical criteria, we benchmarked the model in unconditional generation, conditional generation, and trajectory interpolation.

In unconditional generation, generated cells overlapped real cells in UMAP space for PBMC68k, Tabula Muris Senis and disease-associated lung fibrosis datasets (Fig. 1h,i,j). Generated cells occupied the major regions of the observed transcriptomic manifolds without forming isolated synthetic clusters, indicating that scJET recovers the broad geometry of heterogeneous cell populations. Quantitatively, on PBMC68k, scJET achieved the lowest MMD and Wasserstein-2 distance among the compared methods, while maintaining strong kNN macro-F1 and high gene mean and gene variance correlations (Extended Data Fig. 1c). These results indicate that scJET preserves both distributional structure and gene expression statistics in full-gene expression space.

In conditional generation, scJET reproduced annotated cell classes in Tabula Muris Senis, including immune, epithelial, stromal, endothelial, hepatic, and renal populations (Fig. 2a,b). In PBMCs, it preserved fine immune identities such as monocytes, T cells, B cells, NK cells, dendritic cells, and Tregs (Fig. 2d). At the global distribution level, scJET also achieved superior performance across MMD, Wasserstein distance, kNN accuracy and AUC on the PBMC68k dataset (Fig. 2c), indicating strong alignment between generated and real cellular distributions. Nearest-neighbor and membership inference tests showed no evidence of memorization (Extended Data Fig. 2d–f).

**Fig. 2.**
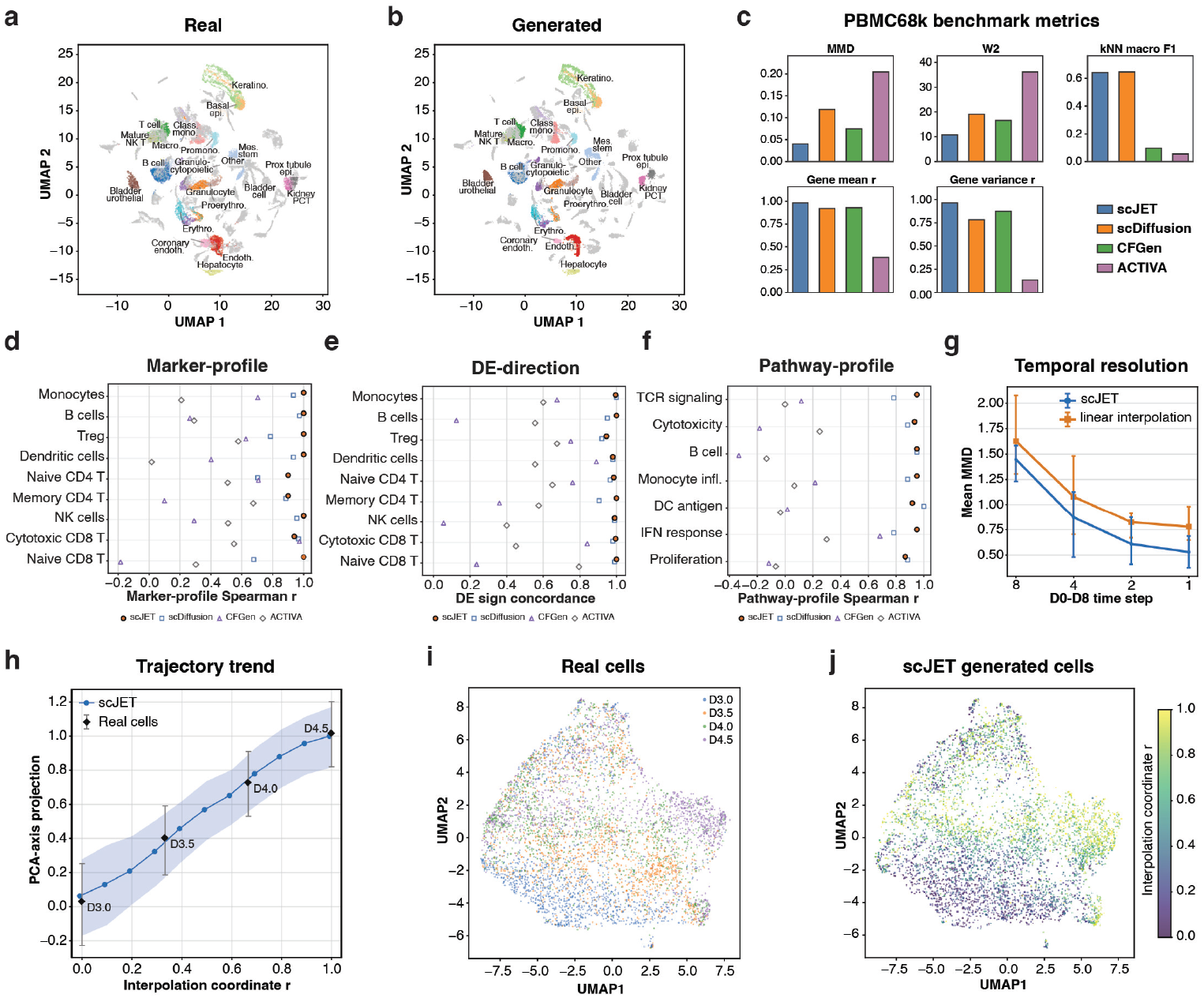
Conditional generation, biological fidelity and trajectory modeling with scJET. (a) UMAP of real cells in Tabula Muris Senis. (b) UMAP of scJET-generated cells. (c) PBMC68k benchmark including MMD, Wasserstein distance, and kNN classification performance. (d) Cell-type marker gene profile fidelity across immune populations. (e) Differential expression directionality consistency across cell types. (f) Pathway-level expression profile consistency across immune programs. (g) Temporal resolution analysis under trajectory interpolation. (h) Trajectory trend along PCA-derived progression axis. (i) UMAP visualization of real cells across temporal stages. (j) UMAP visualization of scJET-generated trajectory manifold.r

To assess biological fidelity beyond global manifold alignment, we examined marker programs, differential-expression directionality, and pathway activity. Even without conditioning, scJET recovered major immune populations and preserved transcriptional programs (Extended Data Fig. 2a–c). scJET achieved consistently high marker gene–expression correlations across cell types, maintained near-perfect differential-expression sign concordance, and preserved pathway-level activity patterns across immune programs including TCR signaling, cytotoxicity, B-cell programs, monocyte inflammation, dendritic antigen presentation, interferon response and proliferation (Fig. 2d-f, Extended Data Fig. 1d-f). Averaged across evaluated categories, scJET achieved a marker gene–expression correlation of 0.971, a differential-expression sign concordance of 0.988 and a pathway activity correlation of 0.931 with real cells (Extended Data Fig. 1f). These results indicate that scJET preserves interpretable transcriptional programs at both gene and pathway levels.

Finally, we evaluated scJET under trajectory interpolation, where intermediate full-gene expression states are generated between time-labeled cellular states using a continuous interpolation coordinate *r*. Across D0-D8 temporal resolutions, scJET produced lower mean MMD than linear interpolation at 8-day, 4-day, 2-day and 1-day intervals, with both methods improving as the temporal step became finer (Fig. 2g). In a D3.0-D4.5 transition segment, scJET followed the monotonic PCA-axis trend of real cells and generated intermediate profiles that aligned with the expected progression pattern (Fig. 2h). UMAP visualization further showed that generated cells formed a continuous *r*-dependent transition over a manifold geometry similar to that of the observed time-labeled cells (Fig. 2i,j). Thus, scJET captures not only static single-cell expression distributions but also continuous transitions within the gene-expression manifold.

scJET establishes expression-space denoising as a scalable strategy for single-cell generative modeling. Unlike VAE, adversarial, and latent-diffusion frameworks that rely on HVG selection, latent compression, or decoder-based reconstruction, scJET operates directly in full gene space through patch-based Transformer denoising. This preserves both high- and low-variance genes while maintaining global manifold structure, local neighborhood statistics, and pathway structure. It supports unconditional generation, cell-type conditional generation, and trajectory interpolation within a unified framework. Importantly, patching is a computational abstraction rather than a biological assumption, enabling efficiency without compromising biological fidelity. The observation that different patch combinations have only a modest effect on generation performance suggests that the patch-based algorithm maintains robust generative capacity while reducing peak computational demand, thereby improving computational efficiency. Overall, scJET provides a basis for AIVC-oriented modeling by preserving transcriptome-wide structure without HVG filtering or latent compression. All datasets are publicly available from published sources, including PBMC68k[18], Tabula Muris Senis[19], idiopathic pulmonary fibrosis (IPF) lung datasets[20], and Waddington-OT lineage tracing datasets [21]. Code used in this study is available at https://github.com/Liangqt-lab/scJET.

## Methods

### Dataset acquisition and cohort definition

We assembled a multi-cohort benchmarking collection of publicly available single-cell RNA-sequencing datasets, spanning peripheral blood mononuclear cells (PBMC68k), Tabula Muris Senis (TMS), idiopathic pulmonary fibrosis (IPF) lung tissues, as well as multiple developmental and perturbation systems with lineage-resolved or time-resolved annotations, including Waddington-OT-derived trajectories when available. All datasets were obtained from processed count matrices reported in the original studies and distributed through GEO or ArrayExpress, and were subsequently harmonized into a unified AnnData-compatible format to ensure consistent downstream processing.

Across all cohorts, the expression data are represented as a matrix *X* ∈ ℝ^*G×C*^ , where *G* denotes the number of genes and *C* the number of cells. In PBMC68k, both a full-gene configuration (*G* = 32,738) and a benchmark subset (*G* = 17,786) were considered, with no highly variable gene filtering applied in the full-gene setting in order to preserve global transcriptomic structure.

Quality control was performed jointly at the cellular and gene levels to ensure robustness of downstream statistical estimation. Cells with fewer than 200 detected genes were excluded, as were genes expressed in fewer than 0.1% of cells or exhibiting zero variance across the full dataset. These criteria were applied uniformly across cohorts, yielding a shared expression space that remains stable across heterogeneous sequencing platforms.

To avoid information leakage across biological conditions, dataset partitioning was defined in a cohort-specific manner. When cell-type annotations were available, splits were stratified to preserve marginal distributions of annotated identities. In lineage-resolved or temporal systems, partitions were instead organized along developmental or progression axes, ensuring that intermediate states were not simultaneously present across training and evaluation subsets. All partitions were fixed prior to model development to ensure full reproducibility.

### Data preprocessing

Raw UMI counts *C*_*ij*_ were first adjusted for sequencing depth variation at the single-cell level. For each cell *i*, total counts are denoted by 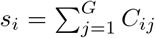, and normalized expression is computed through a log-scaled library-size transformation:

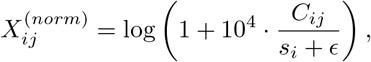

with *ϵ* = 1 introduced to stabilize low-count regimes.

Because scRNA-seq data exhibit heavy-tailed gene-specific expression distributions, we further applied a robust truncation based on empirical quantiles. For each gene *j*, values above the 99.5th percentile 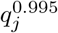 were clipped, yielding 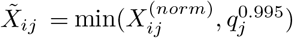. This step preserves global structure while reducing sensitivity to rare transcriptional bursts.

The resulting expression matrix was standardized in a gene-wise manner to align dynamic ranges across genes. For each gene, mean and variance are computed as *µ*_*j*_ and *σ*_*j*_, with a lower bound imposed on *σ*_*j*_ to prevent instability in near-constant genes. Standardized expression is given by

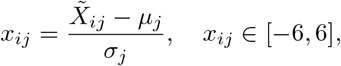

and is subsequently clipped to enforce bounded support. No dimensionality reduction, feature selection, or latent encoding is applied at any stage.

### Full-gene expression representation

Each cell is represented directly in full-gene space as a vector *x*_*i*_ ∈ ℝ^*G*^. To make the representation compatible with Transformer-based sequence modeling, the gene axis is partitioned into contiguous segments of fixed size *P* , yielding a collection of patches 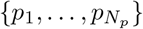 with *N*_*p*_ = ⌈*G/P*⌉ . When the gene dimension is not divisible by *P* , zero-padding is applied to maintain structural consistency.

Each patch is then mapped into a continuous embedding space through a linear projection,

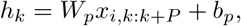

where *W*^*p*^ ∈ ℝ_*P×d*_ produces a *d*-dimensional token representation. Importantly, this patch construction should be interpreted as a computational partitioning of the transcriptome rather than a biologically meaningful segmentation. Gene ordering is fixed by default, although alternative permutations and correlation-informed reorderings are used in ablation analyses to assess robustness to local structure assumptions.

### Model architecture

scJET is formulated as a Transformer-based denoising model operating over patch-level token sequences. Each token integrates patch content with two contextual signals: a continuous corruption-time embedding and an optional conditional embedding encoding biological context. The resulting representation takes the form *z*_*k*_ = *h*_*k*_ + *e*(*t*) + *e*(*y*), where *e*(*t*) encodes the diffusion time and *e*(*y*) represents optional conditioning such as cell identity or experimental state.

The Transformer, parameterized as scJET_*θ*_, maps the token sequence 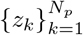 back into the full-gene expression space, producing a reconstruction 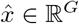. The architecture consists of an 8-layer Pre-LayerNorm Transformer with hidden dimension 480, 10 attention heads, and a feed-forward expansion of 1920 dimensions, using GELU activations throughout. Dropout is not employed in order to preserve deterministic modeling of gene dependencies in the full-gene regime. Conditional control is implemented via classifier-free guidance, where conditioning inputs are randomly replaced by a null embedding during training to support both conditional and unconditional generation at inference.

### Stochastic corruption process

We model single-cell generation as a continuous denoising process in which clean expression states are progressively corrupted by isotropic noise. For a given expression vector *x*, the corrupted observation is constructed as a convex combination:

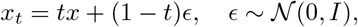

where *t* ∈ (0, 1) controls the corruption level.

Rather than sampling *t* uniformly, we introduce a biased temporal distribution by transforming a Gaussian variable, *t* = *σ*(*η*) with *η* ∼ *N*(− 1, 0.8^2^). This choice concentrates training on intermediate noise regimes, where structural information transitions from local gene-level coherence to global manifold structure. Under this formulation, clean expression profiles can be interpreted as low-dimensional structures embedded within a high-dimensional corrupted space.

### Denoising objective

The model is trained to recover clean expression states from corrupted observations. Given an input *x*_*t*_, the network produces a denoised estimate 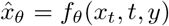, which directly parameterizes the clean expression space.

To stabilize learning across corruption levels, we introduce an auxiliary velocity field,

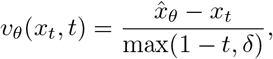

with *δ* = 0.03 preventing divergence in low-noise regimes. The final objective couples reconstruction fidelity with velocity consistency,

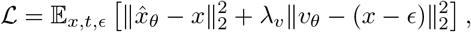

where *λ*_*v*_ = 0.1. This formulation avoids likelihood-based modeling and adversarial training, operating entirely in continuous expression space.

### Sampling procedure

At inference time, scJET generates samples by numerically integrating the learned velocity field starting from isotropic Gaussian noise *x*_0_ ∼ *N*(0, *I*). The dynamics are solved over *t* ∈ [0, 1] using a Heun predictor-corrector scheme with 160 integration steps.

A forward Euler prediction is first computed as

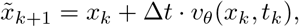

followed by a correction step that symmetrizes velocity evaluations,

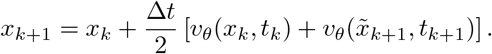

The final state at *t* ≈ 1 is mapped back to expression space using the inverse of the gene-wise standardization statistics learned during preprocessing.

### Training protocol

Model optimization is performed using AdamW with a learning rate of 5 × 10^−5^ and weight decay of 10^−4^. Training uses a batch size of 64, with gradient clipping at a threshold of 1.0 and an exponential moving average (EMA) decay of 0.9995 applied to model parameters. Each model is trained for up to 70,000 optimization steps.

All updates are computed through backpropagation over the denoising objective, and mixed-precision training (FP16) is used to improve computational efficiency while preserving numerical stability. Unless otherwise stated, all hyperparameters remain fixed across datasets to ensure comparability.

### Evaluation metrics

Model performance is evaluated using complementary distributional, discriminative, and biological criteria. Distributional similarity is quantified using Maximum Mean Discrepancy (MMD) with an RBF kernel, where kernel bandwidth is selected using the median heuristic. In addition, Wasserstein-2 distance is computed in a PCA-reduced space (typically 30–50 components), providing a geometric measure of alignment between real and generated distributions.

Discriminative evaluation is performed by training classifiers to distinguish real from generated cells. This includes k-nearest neighbor classifiers (k=5 or 15) and Random Forest models with 300 trees, evaluated under stratified cross-validation. Performance is reported using accuracy, macro-F1, and PRDC-based measures capturing precision, recall, density, and coverage in embedding space.

Biological fidelity is assessed at multiple resolutions. Gene-level agreement is quantified through Pearson and Spearman correlations of gene-wise mean and variance profiles. Marker fidelity is evaluated by comparing canonical cell-type programs, while differential expression consistency is measured as the fraction of preserved log-fold-change signs between real and generated data. At the pathway level, we compute correlations of immune and tissue-specific gene set activities, capturing higher-order biological structure beyond individual gene behavior.

### Ablation studies

We systematically evaluate the contribution of modeling choices across representation, ordering, and architecture. Patch size is varied across *P* ∈ {32, 64, 128} to study the trade-off between token granularity and global context aggregation. Gene ordering is perturbed using default ordering, random permutations, and correlation-informed clustering, allowing assessment of sensitivity to local adjacency assumptions.

Architectural ablations remove or modify key components of the model, including attention layers, conditional embedding pathways, and patch-level interaction mechanisms. In addition, Transformer-based modeling is compared against MLP-based per-patch encoders to isolate the effect of global token mixing. Finally, full-gene scJET is contrasted with latent-variable baselines, including VAE-based denoising models, to quantify the impact of explicit high-dimensional modeling versus compressed latent representations. All comparisons are performed under identical preprocessing and training configurations.

### Trajectory modeling

To model continuous transitions in cellular state space, we introduce a scalar interpolation variable *r* ∈ [0, 1] defined between endpoint populations derived from Waddington-OT or equivalent lineage reconstructions. Intermediate states are generated by conditioning scJET on this progression variable, producing continuous trajectories in full-gene expression space.

Trajectory quality is assessed through multiple complementary criteria. We measure distributional agreement with real intermediate states using MMD, and evaluate geometric consistency along principal components associated with temporal variation. In addition, smoothness is quantified via cumulative displacement along the trajectory, ensuring that generated sequences preserve coherent biological transitions while avoiding discontinuities in expression space.

## Acknowledgments

Q.R.L. and Q.T.L. designed the study. Q.T.L. performed all biostatistical analyses and modeling and drafted the manuscript. Q.R.L. supervised the study and revised the manuscript.

---

**Extended data Fig. 1.**
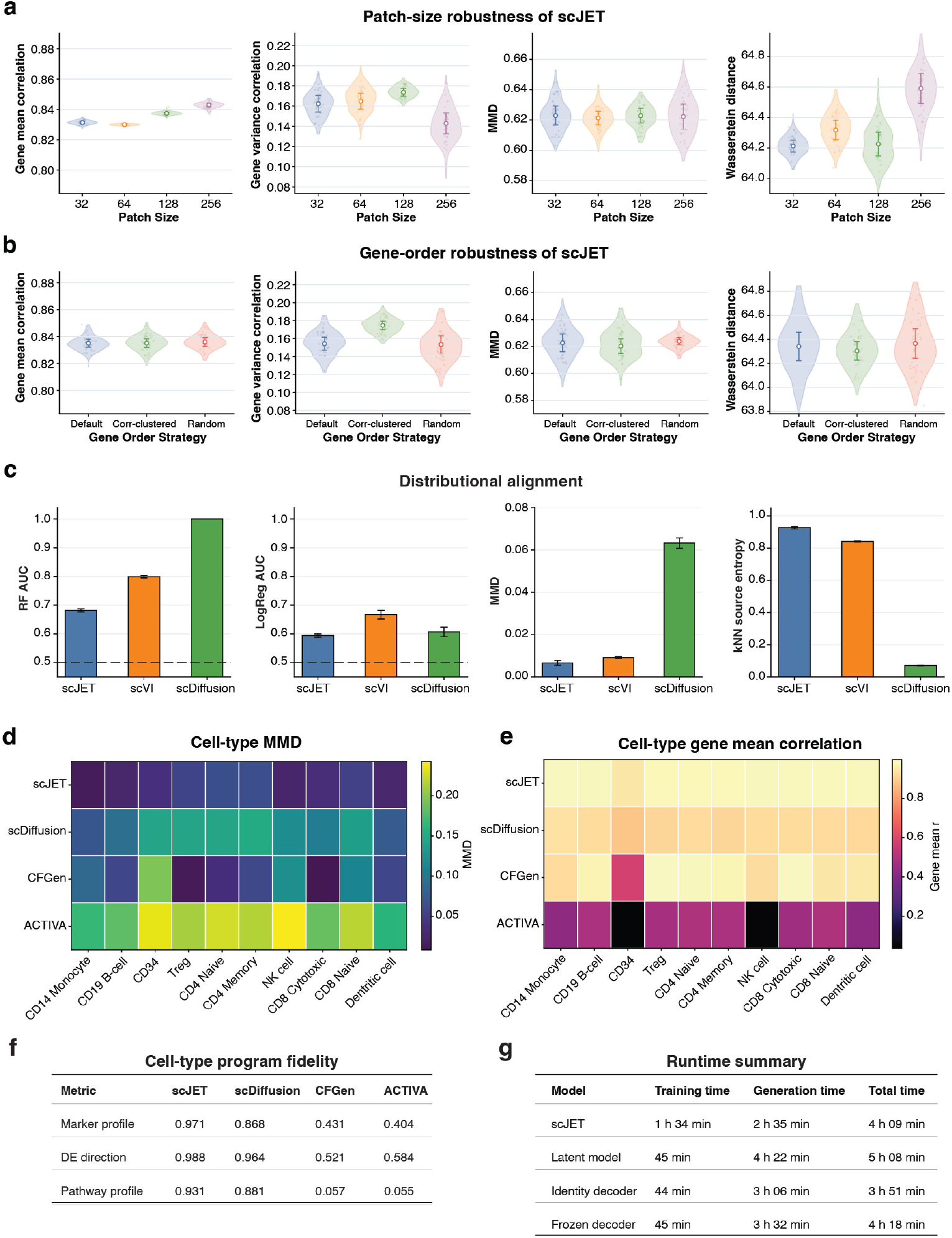
Evaluation of scJET. (a) Patch-size robustness analysis. (b) Gene-order robustness under different permutation strategies. (c) Benchmarking across scJET, scVI, scdiffusion. (d) Cell-type MMD across methods. (e) Cell-type gene mean correlation heatmap across methods. (f) Cell-type program fidelity across marker profiles, differential expression directionality, and pathway activity. (g) Runtime and computational efficiency comparison across models.

**Extended data Fig. 2.**
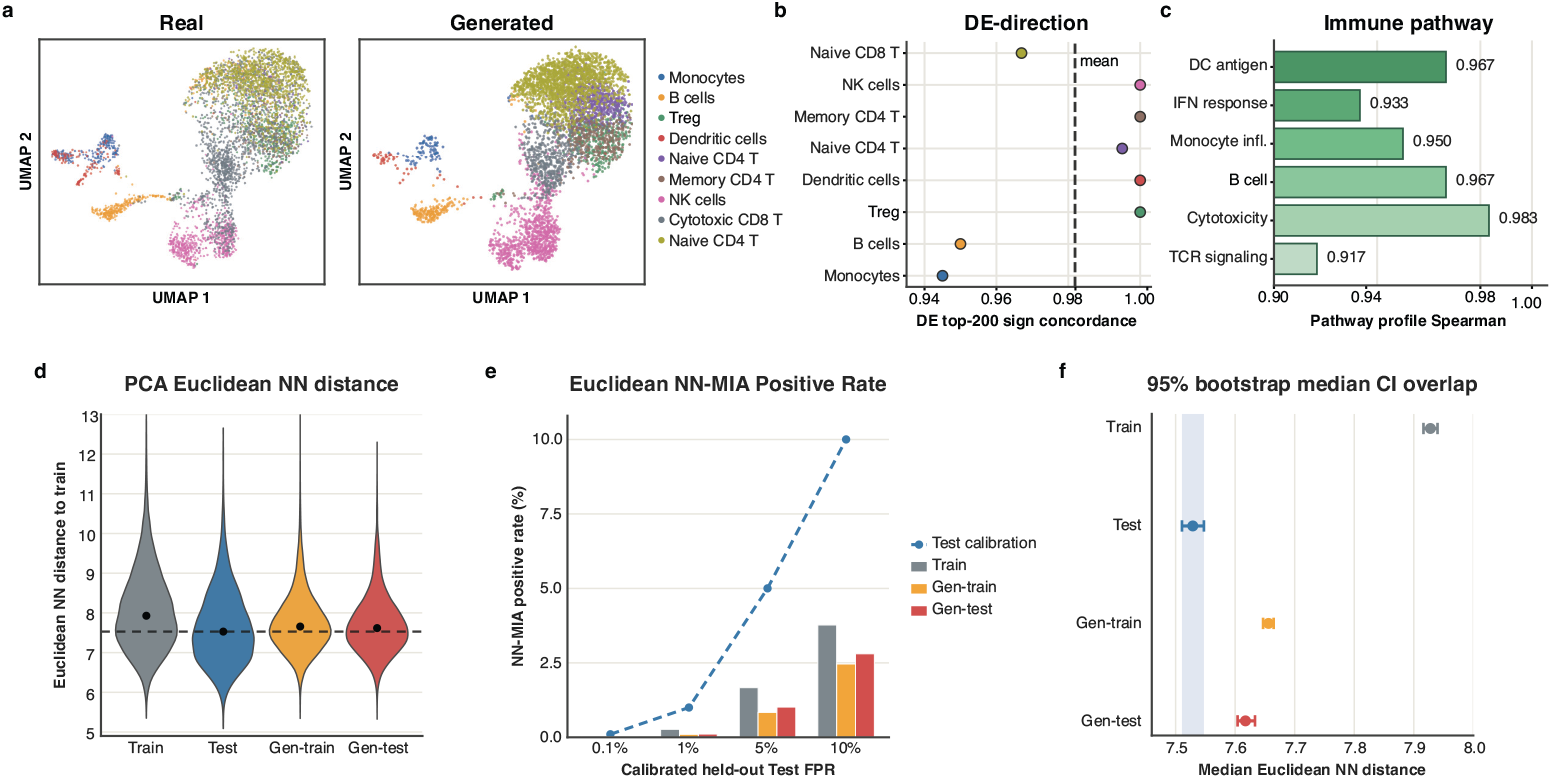
Biological fidelity and memorization analysis of scJET on PBMC68k. (a) UMAP of real and scJET-generated PBMC68k cells. (b) Differential expression directionality consistency across cell types. (c) Immune pathway activity correlation between real and generated cells across cell types. (d) PCA-space Euclidean nearest-neighbor(NN) distance distributions between real, held-out test, and generated cells (Gen-train: generation conditioned on training-derived prior; Gen-test: generation conditioned on test-matched prior). (e) NN-based membership inference positive rates under calibrated false-positive thresholds. (f) Bootstrap analysis of median nearest-neighbor distances with 95% confidence intervals (3,000 resamples).

## References

[1] Bunne, C. et al. How to build the virtual cell with artificial intelligence: priorities and opportunities. Cell 187, 7045–7063 (2024).

[2] Yang, T., Wang, Y. Y., Ma, F., Xu, B. H. & Qian, H. L. Build the virtual cell with artificial intelligence: a perspective for cancer research. Military Medical Research 12, 4 (2025)

[3] Dibaeinia, P. et al. Virtual cells need context, not just scale. bioRxiv (2026). Preprint.

[4] Lopez, R., Regier, J., Cole, M. B., Jordan, M. I. & Yosef, N. Deep generative modeling for single-cell transcriptomics. Nature Methods 15, 1053–1058 (2018).

[5] Marouf, M. et al. Realistic in silico generation and augmentation of single-cell RNA-seq data using generative adversarial networks. Nature Communications 11, 166 (2020).

[6] Heydari, A. A., Davalos, O. A., Zhao, L., Hoyer, K. K. & Sindi, S. S. Activa: realistic single-cell rna-seq generation with automatic cell-type identification using introspective variational autoencoders. Bioinformatics 38, 2194–2201 (2022).

[7] Song, D. et al. scDesign3 generates realistic in silico data for multimodal single-cell and spatial omics. Nature Biotechnology 42, 247–252 (2024).

[8] Zappia, L., Phipson, B. & Oshlack, A. Splatter: simulation of single-cell rna sequencing data. Genome Biology 18, 174 (2017).

[9] Luecken, M. D. & Theis, F. J. Current best practices in single-cell rna-seq analysis: a tutorial. Molecular systems biology 15, MSB188746 (2019).

[10] Hicks, S. C., Townes, F. W., Teng, M. & Irizarry, R. A. Missing data and technical variability in single-cell rna-sequencing experiments. Biostatistics 19, 562–578 (2018).

[11] Lähnemann, D. et al. Eleven grand challenges in single-cell data science. Genome biology 21, 31 (2020).

[12] Stegle, O., Teichmann, S. A. & Marioni, J. C. Computational and analytical challenges in single-cell transcriptomics. Nature Reviews Genetics 16, 133–145 (2015).

[13] Ho, J., Jain, A. & Abbeel, P. Denoising diffusion probabilistic models, Vol. 33, 6840–6851 (2020).

[14] Luo, E., Hao, M., Wei, L. & Zhang, X. scDiffusion: conditional generation of high-quality single-cell data using diffusion model. Bioinformatics 40, btae518 (2024).

[15] Zhang, T. et al. cfDiffusion: diffusion-based efficient generation of high quality scRNA-seq data with classifier-free guidance. Briefings in Bioinformatics 26, bbaf071 (2025).

[16] Li, T. & He, K. Back to basics: Let denoising generative models denoise (2026). ArXiv:2511.13720v2, arXiv:2511.13720.

[17] Moon, K. R. et al. Visualizing structure and transitions in high-dimensional biological data. Nature biotechnology 37, 1482–1492 (2019).

[18] Zheng, G. X. et al. Massively parallel digital transcriptional profiling of single cells. Nature communications 8, 14049 (2017).

[19] A single-cell transcriptomic atlas characterizes ageing tissues in the mouse. Nature 583, 590–595 (2020).

[20] Adams, T. S. et al. Single-cell rna-seq reveals ectopic and aberrant lung-resident cell populations in idiopathic pulmonary fibrosis. Science advances 6, eaba1983 (2020).

[21] Schiebinger, G. et al. Optimal-transport analysis of single-cell gene expression identifies developmental trajectories in reprogramming. Cell 176, 928–943 (2019).

